# Mutated in Colorectal Cancer (MCC) is a centrosomal protein that relocalizes to the ncMTOC during intestinal cell differentiation

**DOI:** 10.1101/2021.07.27.453941

**Authors:** Lucian B. Tomaz, Bernard A. Liu, Sheena L. M. Ong, Ee Kim Tan, M. Meroshini, Nicholas S. Tolwinski, Christopher S. Williams, Anne-Claude Gingras, Marc Leushacke, N. Ray Dunn

## Abstract

*Mutated in Colorectal Cancer (MCC)* encodes a coiled-coil protein implicated, as its name suggests, in the pathogenesis of hereditary human colon cancer. To date, however, the contributions of MCC to intestinal homeostasis remain unclear. Here, we examine the subcellular localization of MCC, both at the mRNA and protein levels, in the adult intestinal epithelium. Our findings reveal that Mcc transcripts are restricted to proliferating crypt cells, including *Lgr5+* stem cells, and that Mcc protein is distinctly associated with the centrosome in these cells. Upon intestinal cellular differentiation, Mcc is redeployed to the non-centrosomal microtubule organizing center (ncMTOC) at the apical domain of villus cells. Using intestinal organoids, we show that the shuttling of the Mcc protein depends on phosphorylation by Casein Kinases 1δ/ε, which are critical modulators of WNT signaling. Together, our findings support a putative role for MCC in establishing and maintaining the cellular architecture of the intestinal epithelium as a component of both the centrosome and ncMTOC.

## Introduction

*Mutated in Colorectal Cancer (MCC)* was identified through cytogenetic and linkage studies as a culprit tumor suppressor gene for the autosomal dominant human hereditary colon cancer syndrome Familial Adenomatous Polyposis (FAP) (Kinzler et al., 1991b; a). FAP patients typically present hundreds to thousands of adenomas in the colon and rectum (Waller et al., 2016). Later the same year, *Adenomatous Polyposis Coli (APC)*, which is tightly linked to *MCC* on human chromosome 5q21, was correctly established as the gene responsible for FAP (Groden et al., 1991; Kinzler et al., 1991a; Nishisho et al., 1991). Despite its historical association with colorectal cancer (CRC), linkage to *APC* and strong evolutionary conservation (Luongo et al., 1993), *MCC* transcript distribution, the subcellular localization of the MCC protein and its precise function in the intestine have not been fully characterized.

The *MCC* gene encodes a large coiled-coil protein harboring a highly conserved, extreme C-terminal type I (PSD-95/Dlg/ZO-1) PDZ binding motif (PBM) (Bourne, 1991; Arnaud et al., 2009; Pangon et al., 2012). Depending on the cell line, tissue, reagent or assay used, MCC has been found in several organelles (mitochondria and endoplasmic reticulum) and cellular compartments (plasma membrane, cytoplasm and nucleus) (Arnaud et al., 2009; Benthani et al., 2018; Fukuyama et al., 2008; Senda et al., 1997; Matsumine et al., 1996; Pangon et al., 2010). These discrepant results have casted doubt about the precise role that MCC serves within a cell. For example, MCC is known to be a phosphoprotein, with phosphorylation at position −1 required for its interaction with the PDZ-domain protein SCRIBBLE at the active migratory edge of CRC cell lines (Pangon et al., 2012; Caria et al., 2019). When overexpressed, MCC has been shown to bind β-catenin (CTNNB1) in the nucleus to negatively regulate canonical WNT signaling in cancer cell lines and to inhibit cell proliferation (Fukuyama et al., 2008; Pangon et al., 2015). In contrast, a recent report argues that MCC is membrane associated and interacts with β-catenin and E-cadherin to strengthen cell adhesion in CRC cell lines (Benthani et al., 2018). These confounding results motivated us to identify the MCC protein interactome in an effort to firmly define its subcellular localization and function.

Here, we show for the first time that MCC associates with the protein interaction network that surrounds the centrosome in proliferating cells, known as the major microtubule organizing center (MTOC) (Brinkley, 1985; Muroyama and Lechler, 2017), both in assorted cell lines and within mouse and human intestinal crypts. Upon exit from the cell cycle and terminal differentiation, we find that centrosomal MCC protein redeploys from the MTOC to the apical membrane of enterocytes, incorporating into the non-centrosomal microtubule organizing center (ncMTOC) that anchors the minus end of microtubules and establishes apicobasal polarity (Meads and Schroer, 1995; Goldspink et al., 2017b; Muroyama and Lechler, 2017). Lastly, we provide evidence that the relocalization of the Mcc protein from the MTOC to the ncMTOC is governed by Casein Kinase 1δ/ε phosphorylation, whose interaction with MCC was revealed by our proteomics studies and implicates WNT signaling in this crucial process that sustains intestinal homeostasis.

## Results

### *Mcc* is specifically expressed in crypts of the intestinal epithelium

The intestinal epithelium is a constantly renewing single cell layer organized into crypt and villus units (Fig. *1A*) (Gehart and Clevers, 2019). Finger-like villi harboring differentiated epithelial cells project into the intestinal lumen to facilitate nutrient absorption. Each villus is encircled by multiple contiguous, proliferative crypt compartments embedded within the underlying submucosa that contain crypt base columnar (CBC) stem cells (Leushacke and Barker, 2014; Barker, 2014; Clevers, 2013). We previously reported using an *Mcc^lacZ^* reporter allele that *Mcc* is expressed in the adult mouse intestine (Young et al., 2011). However, the specific cell populations expressing *Mcc* in the crypt and villus were not fully characterized. We therefore re-examined *Mcc^lacZ^* expression by detection of β-Galactosidase (β-Gal) activity using meticulously controlled 5-bromo-4-chloro-3-indolyl-β-d-galactopyranoside/ferri-/ferrocyanide (X-Gal) histochemistry (Merkwitz et al., 2016). We found that β-Gal activity is restricted to crypt units in both the small intestine (Fig. *1B, C*) and colon (Fig. *1D*). Co-staining for β-Gal and Intestinal Alkaline Phosphatase (IAP), a marker for differentiated villus cells (Sussman et al., 1989; Hinnebusch et al., 2004), in mechanically isolated crypt and villus fractions revealed that β-Gal-positive cells are distributed along the entire crypt, but not in the villus (Supp. *Fig. 1A-B*). Significantly, no detection of β-Gal activity was observed in wholemount wild-type (WT) crypt fractions used as a negative control for our X-Gal histochemical analysis (Supp. Fig. *1B*).

**Figure 1.**
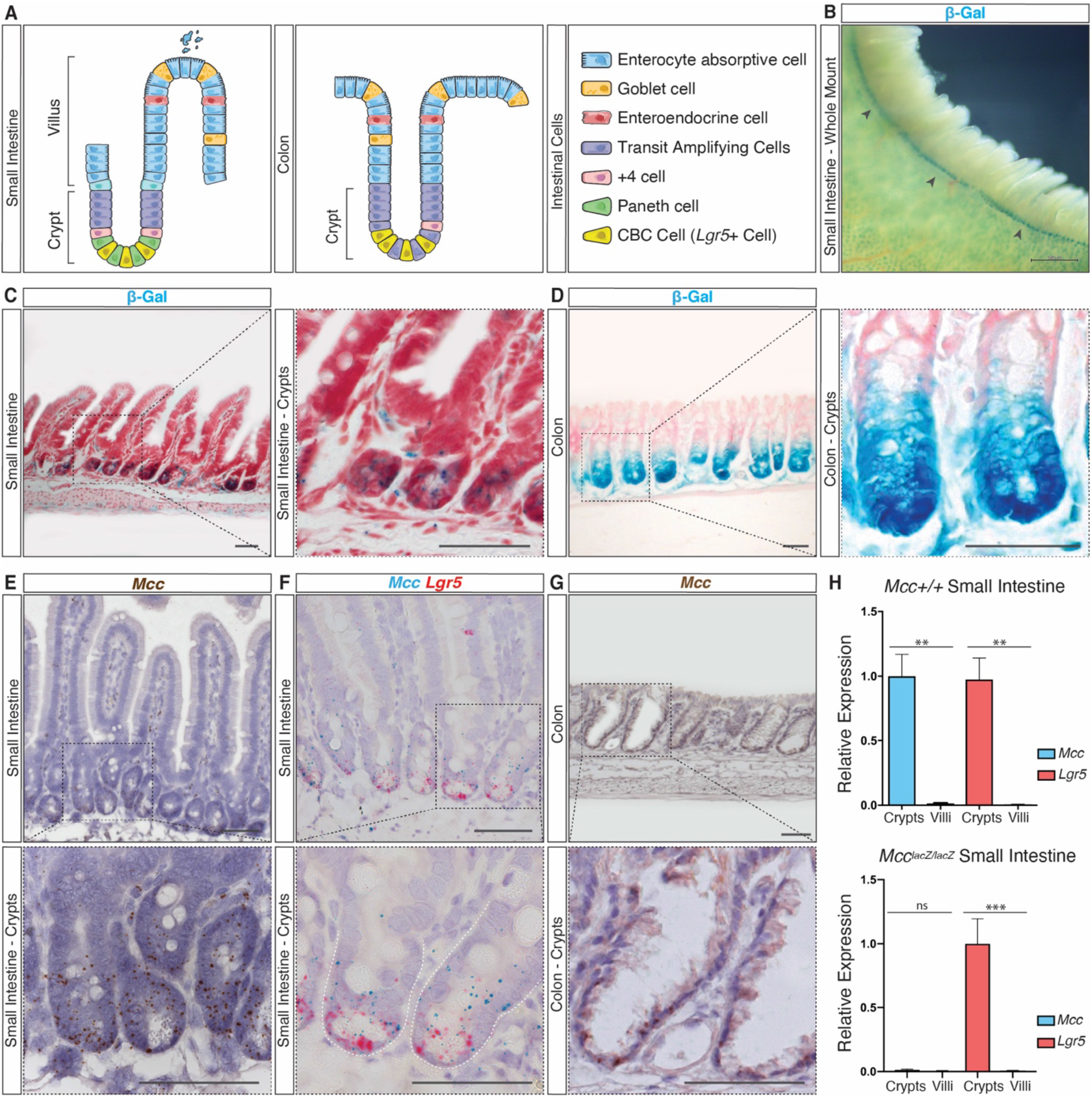
*Mcc* is specifically expressed in crypts of the intestinal epithelium. *(A):* Schematic representation of small intestine (SI) and colon epithelia cellular organization. *(B):* Histochemical (HC) staining for β-galactosidase (β-Gal) activity in whole-mount *Mcc^lacZ/lacZ^* SI. Black arrowheads indicate β-gal activity (blue) restricted to crypt units. Scale bar, 500 μm. *(C-D):* HC for β-Gal activity on sections of *Mcc^lacZ/lacZ^* SI and colon tissues counter-stained with Nuclear Fast Red. Scale bars, 50 μm. *(E-G):* Section *in situ* hybridization (ISH) on wild-type (WT) SI and colon tissues. *(E)* Endogenous *Mcc* expression is specifically observed in crypts. *(F): Mcc* (blue) expression overlaps with *Lgr5* (red) at the crypt base but extends into the transit-amplifying compartment. *(G):* Endogenous *Mcc* expression on WT colonic crypts. Scale bars, 50 μm. Black-dashed squares in *(C-G)* highlight regions selected for higher magnification. *(H):* qPCR analysis for *Mcc* and *Lgr5* in purified crypt and villus fractions from WT and *Mcc^lacZ/lacZ^* SI. Data were tested for significance by an unpaired two-tailed t-test. P-values of statistical significance are represented as ***p < 0.001, **p < 0.01; n=3.

Next, we performed high-resolution RNA section *in situ* hybridization (ISH) for endogenous *Mcc* transcripts singly or in combination with *Leuoine-rioh repeat containing G-protein coupled receptor 5 (Lgr5)*, whose expression specifically labels CBC cells (Barker et al., 2007) (Fig. *1A,E-G*; Supp. Fig. *1C-D*). *Mcc* and *Lgr5* transcripts overlap in CBC cells at the crypt base, while *Mcc* transcripts extend distally into the transit-amplifying (TA) compartment (Fig. *1A,E-F*). Similarly, *Mcc* expression is restricted to crypts of the colonic epithelium (Fig. *1A, G*). We further show that *Mcc* and *Lgr5* are co-expressed in intestinal crypts using quantitative PCR (qPCR) (Fig. *1H*). Taken together, our findings show that *Mcc* expression is entirely absent in differentiated cells of the small intestine and colon epithelia.

### Proteomics analyses reveal MCC as a centrosomal protein

To provide insight into both MCC protein function and subcellular localization in the intestine, we first chose to establish the MCC protein-protein interactome in HEK293 cells which express MCC endogenously (Arnaud et al., 2009). To accomplish this, we overexpressed FLAG-tagged human *MCC* followed by immunoprecipitation with anti-FLAG agarose beads and mass spectrometry (MS) (Supp. Fig. *2A-D*). Bioinformatic studies revealed that MCC interacts with several centrosomal proteins such as CEP131, CEP170 and NDE1; with the casein kinases CSNK1D (CK1δ) and CSNK1E (CK1ε); and with the PDZ-domain containing polarity proteins such as SCRIBBLE (SCRIB), SNX27 and NHERF1/2 (Fig. *2A*). We further confirmed that the interaction between MCC and either SCRIB or NHERF1 (Arnaud et al., 2009; Pangon et al., 2012) was completely dependent on the PBM of MCC (Supp. Fig. 2*E*). Several proteins that regulate small GTPases were also recovered, including RASAL2, IQGAP1 and RAB11FIP5 (also known as RIP11) (Fig. *2A*). These interactions were independently confirmed by immunoprecipitation of Myc-tagged MCC and Western blotting for endogenous proteins, including NDE1, CEP131 and CEP170, SCRIB and CK1ε (Fig. *2B*).

**Figure 2.**
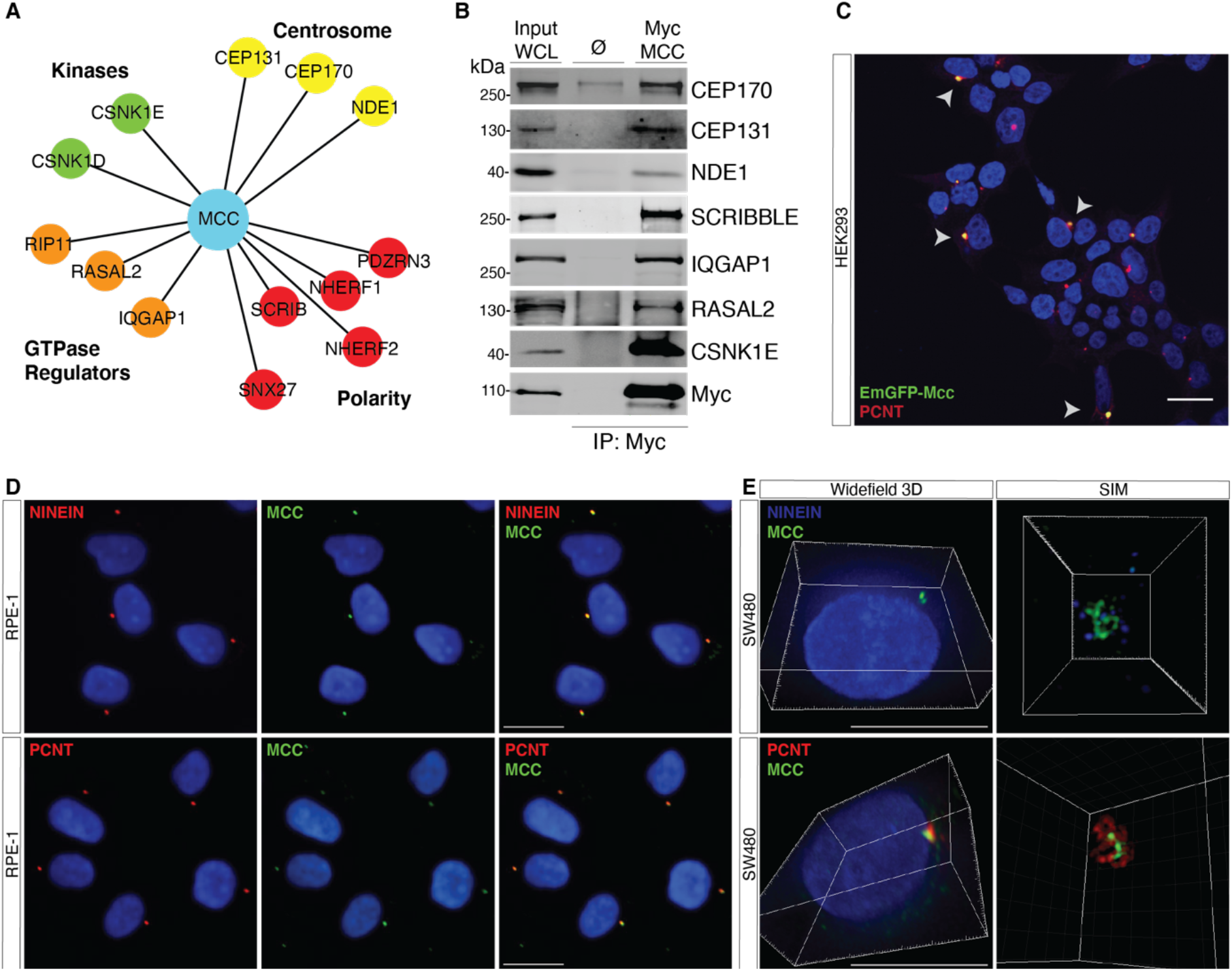
Proteomics analyses reveal MCC as a centrosomal protein. *(A):* Immunoprecipitation (IP) of FLAG-tagged human MCC in HEK293 cells followed by mass spectrometry identifies various MCC interactors, including centrosomal proteins (yellow), cell polarity proteins (red), GTPase regulators (orange), and kinases (CSNK1E/D) (green). *(B):* IP of Myc-tagged human MCC in HEK293 cells followed by Western blotting for indicated interactions. WCL, whole cell lysate. Ø, *pCMV* empty vector control. *(C):* HEK293 cells transfected with *EmGFP-Mcc*. Arrows indicate colocalization of EmGFP-Mcc with the centrosomal protein PERICENTRIN (PCNT). Scale bar, 20 μm. *(D):* Immunofluorescence (IF) for MCC and NINEIN or PCNT shows colocalization at the centrosome in human RPE-1 cells. *(E):* IF 3D widefield showing MCC colocalizing with PCNT and NINEIN. Super resolution Structured Illumination Microscopy (SIM) showing spatial proximity between MCC and PCNT or NINEIN at the centrosome in SW480 cells. Scale bars, 50 μm.

A number of reports have described the subcellular localization of MCC, with often conflicting results (Arnaud et al., 2009; Benthani et al., 2018; Fukuyama et al., 2008; Senda et al., 1997; Matsumine et al., 1996; Pangon et al., 2010). Our interactome studies provided the first indication that MCC might reside at the centrosome. To explore this possibility, we generated an N-terminal *EmGFP*-*Mcc* expression construct and transfected it into HEK293 cells. We observed colocalization of GFP with endogenous PERICENTRIN (PCNT), a component of the Pericentriolar Material (PCM) of the centrosomal complex (Doxsey et al., 1994) (Fig. *2C*). To further characterize MCC localization, we identified a commercially available antibody that shows highly specific staining for endogenous MCC by immunofluorescence (IF) (Table 01). We then performed IF for MCC and PCNT as well as a second centrosomal protein, NINEIN (NIN) (Doxsey et al., 1994; Bouckson-Castaing et al., 1996) in a variety of cell lines, including RPE-1, SW480, HEK293, and HTC116. Microscopy analysis using both confocal and super resolution techniques such as Structured Illumination Microscopy (SIM) (Heintzmann and Huser, 2017) confirmed MCC colocalization with either PCNT or NIN (Fig. *2D, E*; Supp. Fig. *3A*). We also detected colocalization of MCC at the centrosome with several of its interacting partners identified by MS, including CEP170, NDE1, CEP131 and NHERF1 (Supp. Fig. *3B*). Quantitative analysis of numerous IF images in RPE-1 cells revealed that the colocalization of MCC with NIN, CEP170, NDE1, CEP131 and NHERF1 shows a strong positive correlation (*r*) (Supp. Fig. *3C*).

**Table 01.** Primary Antibodies.

| Antibody | Species | Catalog # | Brand | Application |
| --- | --- | --- | --- | --- |
| β-actin | Rabbit | AB8227 | Abcam | WB, IF, IP |
| β-catenin | Rabbit | AB32572 | Abcam | WB, IF |
| β-tubulin | Mouse | T8328 | Sigma | WB, IF, IP |
| Csnk1e | Mouse | (A6) SC-374069 | Santa Cruz | WB, IP |
| Cep131 | Rabbit | PA5-54953 | Invitrogen | WB, IF, IP |
| Cep170 | Rabbit | AB72505 | Abcam | WB, IF, IP |
| Ep-CAM | Rabbit | AB71916 | Abcam | IF |
| E-cadherin | Rabbit | AB15148 | Abcam | IF |
| Iqgap1 | Mouse | (C9) SC-376021 | Santa Cruz | IP |
| Mcc | Mouse | SC-135982 | Santa Cruz | WB, IF, IP |
| Nde1 | Rabbit | PA5-87297 | Invitrogen | WB, IF, IP |
| Ninein | Mouse | 637327 | Merck | IF |
| Ninein | Rabbit | AB4447 | Abcam | IF |
| Nherf1 | Rabbit | SC-271552 | Santa Cruz | WB, IF, IP |
| Pericentrin | Rabbit | AB4448 | Abcam | IF |
| Rasal2 | Rabbit | A302-109A | Bethyl Lab | WB, IF |
| Scribble (H300) | Rabbit | SC-28737 | Santa Cruz | WB, IF, IHC |

In summary, our study using a combination of MS, tagged protein overexpression and extensive IF reveals that MCC is a protein associated with the centrosome. We attribute the difference between our present results and earlier observations to technical factors involving the antigenicity of centrosomal proteins, including the choice of fixation and permeabilization regents, as previously discussed, as well as antibody selection (Hua and Ferland, 2017; Goldspink et al., 2017a).

### Mcc localizes to the centrosome in crypt cells and apical membrane of villus cells

We next performed IF for Mcc in mouse and human intestinal sections. Confocal microscopy confirmed that Mcc specifically localizes to the centrosome in proliferating crypt cells of both the small intestine (Fig. *3A, B*; Supp. Fig. *4A*) and colon (Supp. *Fig 4B*). Strikingly, however, we observed highly specific staining for Mcc at the apical membrane of differentiated cells in the villus compartment (Fig. *3C, D*;Supp. Fig. *4C*) and surface of the colonic epithelium (Supp. *Fig 4B*). No signal for Mcc was detected in crypt and villus sections of homozygous *Mcc* null mice that were used as a negative control for all IF analyses (Supp. Fig. *4D, E*). We additionally examined the presence of Mcc protein in lysates from purified crypt and villus fractions by western blot. Consistent with our IF results, Mcc protein was detected in WT crypt and villus fractions (Fig. *3E*). Moreover, we observed colocalization of Mcc with Pcnt at the centrosome in crypt cells and at the apical membrane in differentiated villus cells (Fig. *3F*; Supp. *Fig. 4F*). Further IF colocalization studies revealed that Mcc overlaps with β-catenin specifically at the apico-lateral junction in both cycling and differentiated intestinal cells (Supp. Fig. *4G, H*). The interaction between MCC and β-catenin *in vitro* has been previously described (Benthani et al., 2018). Collectively, our findings show that Mcc specifically localizes to the centrosome (MTOC) in intestinal crypt cells and as cells undergo terminal differentiation, Mcc is redeployed to the ncMTOC at the apical membrane of differentiated villus cells.

**Figure 3.**
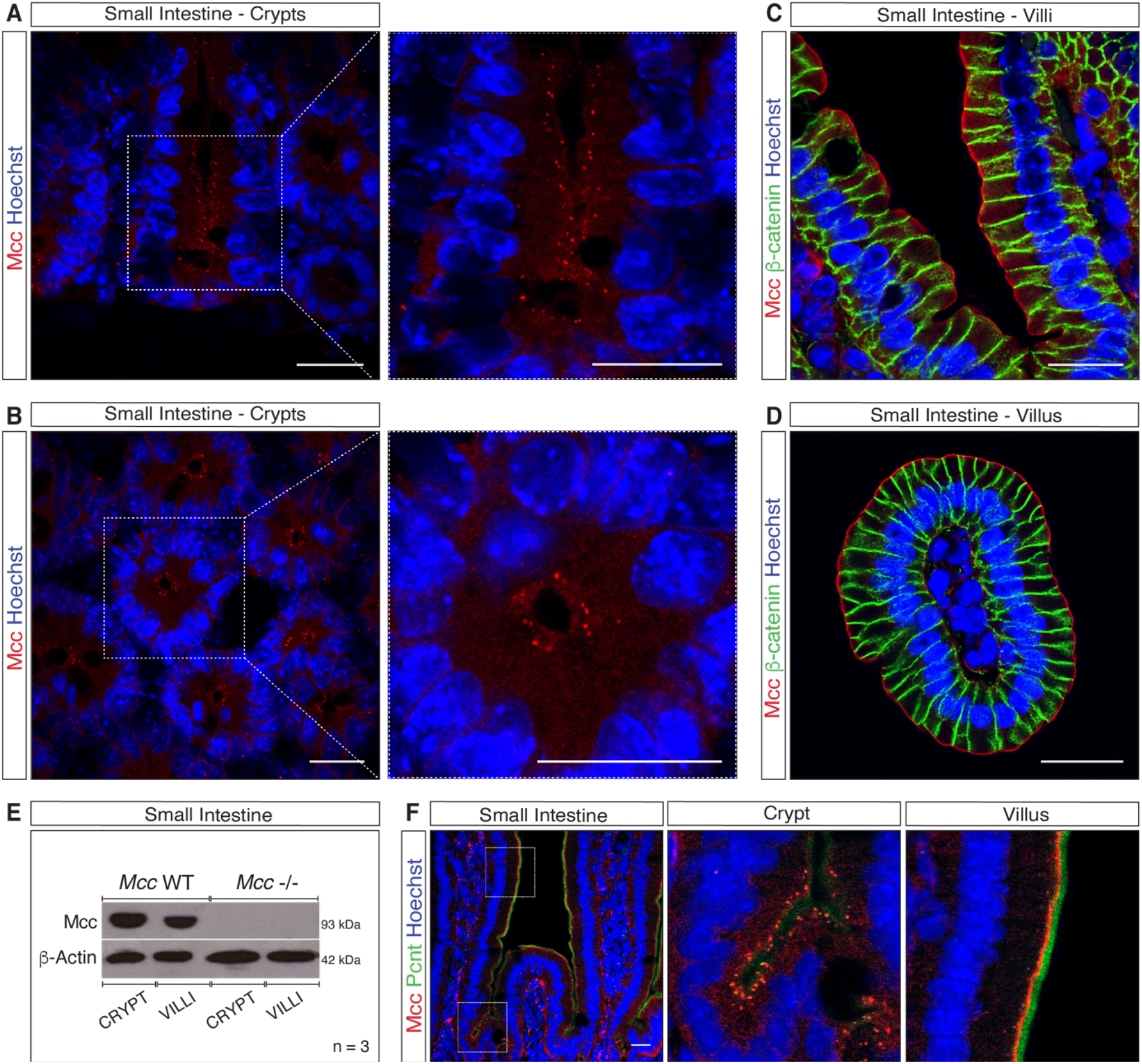
Mcc localizes to the centrosome in crypt cells and apical membrane of villus cells. *(AD):* Immunofluorescence (IF) for Mcc in the mouse small intestine (SI). White-dashed squares highlight regions selected for higher magnification (*A* and *C*, longitudinal and *B* and *D*, transverse sections). *(A)* and *(B):* Punctate centrosomal staining for Mcc is observed in crypt cells. *(C)* and *(D):* IF for Mcc and β-catenin in SI villi. Mcc localizes to the apical membrane while β-catenin labels the lateral membrane of villus cells. Scale bars, 20 μm. *(E):* Western blot for Mcc in whole-cell lysates from purified crypt and villus fractions of the mouse SI. *(F):* IF showing colocalization of Mcc with the centrosomal protein Pericentrin (Pcnt) in the crypt and villus units of the mouse SI. White-dashed squares highlight regions selected for higher magnification. Scale bar, 20 μm.

### Phosphorylation by CK1δ/ε triggers MCC redeployment to the ncMTOC at the apical membrane of villus cells

MCC is a phosphoprotein (Pangon et al., 2012; Caria et al., 2019; Arnaud et al., 2009). Significantly, among the MCC-interacting proteins uncovered by our MS analysis are two Casein Kinases 1 (CK1δ and ε) known to play both positive and negative roles in the WNT/β-Catenin signaling pathway and regulatory functions at the centrosome (Peters et al., 1999; Sakanaka et al., 1999; Gao et al., 2002; Cruciat, 2014; Greer et al., 2014). We therefore asked whether MCC is a direct target of CK1δ/ε phosphorylation. Western blot analysis of SW480 cell lysates confirmed two closely migrating bands of endogenous MCC, one at 93 kDa, which is consistent with the predicted molecular weight of the unmodified protein (Pangon et al., 2010), and a second, higher molecular weight protein that was absent following treatment of the cells with PF670462 (Fig. *4A*), a highly specific inhibitor of CK1δ/ε (Badura et al., 2007; Keenan et al., 2018). This result indicates that MCC is a target of CK1δ/ε phosphorylation. To confirm this finding, we co-expressed Myc-MCC with either FLAG-CK1ε or a catalytically inactive form of CK1ε (FLAG-CK1ε^D128A^) in HEK293 cells. Only the active form of CK1ε induced an upward mobility shift of MCC (Fig. *4B*). We additionally confirmed that overexpression of Myc-MCC and FLAG-CK1ε in the presence of PF670462 also abolishes the higher molecular weight form of MCC, revealing that phosphorylation of MCC is triggered by CK1ε (Fig. *4C*).

**Figure 4.**
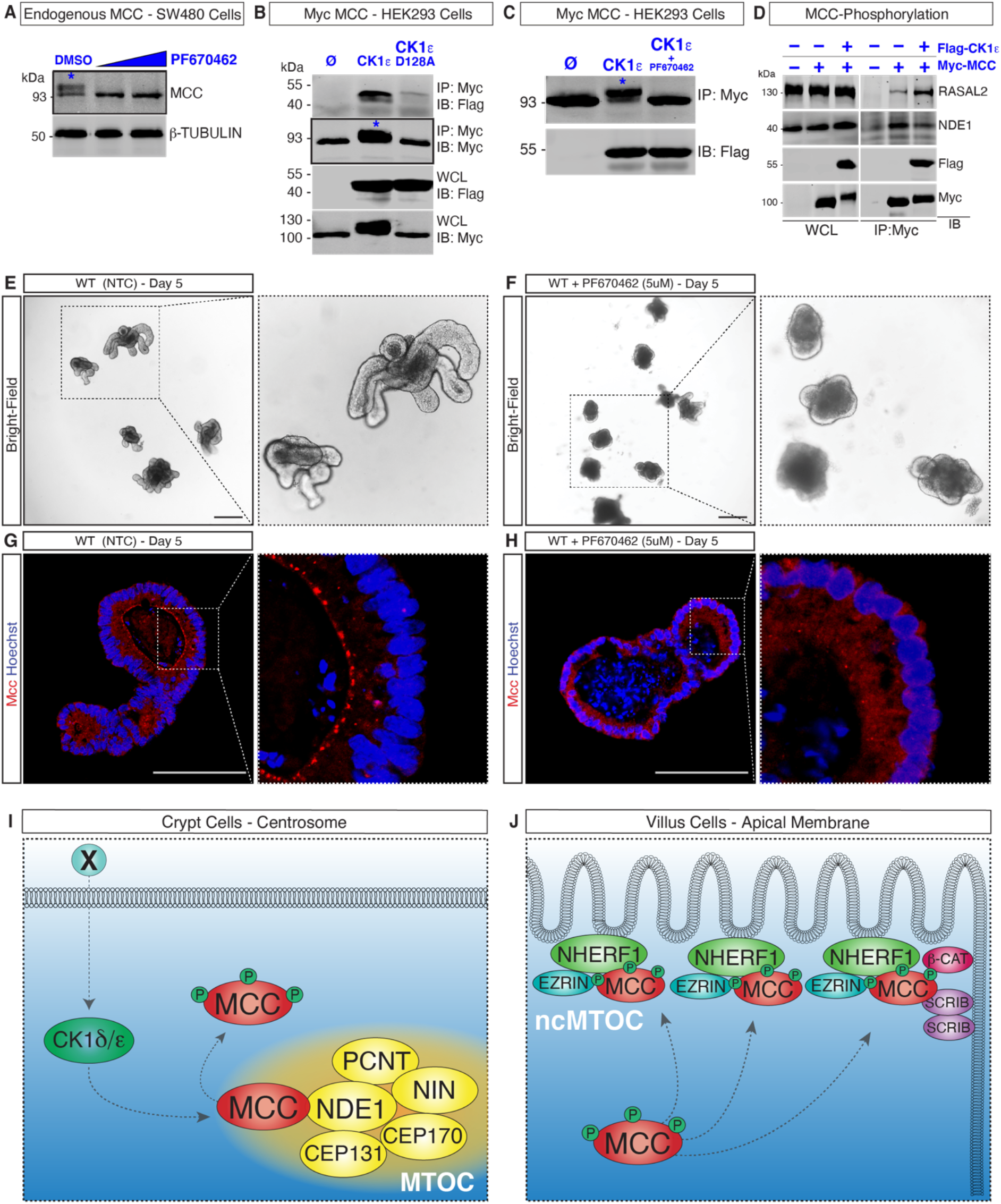
Phosphorylation by CK1δ/ε triggers MCC redeployment to the ncMTOC at the apical membrane of villus cells. *(A):* Western blot for MCC in SW480 colon cancer cells. Treatment with PF67046 (5 μM and 10 μM) eliminates the phosphorylated MCC (asterisk, higher-molecular weight in the DMSO lane). *(B):* Co-expression of Myc-MCC and FLAG-CK1ε in HEK293 cells results in the phosphorylation of Myc-MCC (asterisk, lane 2). Phosphorylation of MCC is not observed with CK1ε-D128A (catalytically inactive). Ø = Empty. *(C):* Treatment with PF670462 eliminates the phosphorylated form of Myc-MCC (upper band, asterisk) in HEK293 cells co-transfected with FLAG-CK1ε. *(D):* Coexpression of Myc-MCC and FLAG-CK1ε in HEK293 cells followed by immunoblotting (IB) analysis reveal that the interaction of MCC with RASAL2 is stabilized upon MCC phosphorylation, while the interaction with NDE1 is weakened. *(E-F):* Wholemount widefield images of mouse wild-type (WT) small-intestinal organoids and *(F)* WT organoids treated with 5 μM of PF670462. *(G):*Immunofluorescence (IF) showing Mcc localization along the apical membrane of differentiated cells in WT organoids. *(H):* Mcc localization at the apical membrane is disrupted upon treatment with 5 μM of PF670462 in WT organoids. White-dashed squares highlight regions selected for higher magnification. Scale bars, 50 μm. *(I-J):* Working model for MCC redeployment during intestinal cell differentiation. *(I):*Phosphorylation of MCC by CK1δ/ε releases MCC from the MTOC by decreasing its affinity to centrosomal proteins such as NDE1. *(J):* MCC relocalizes to the ncMTOC interacting with NHERF1 at the apical membrane, and with SCRIB and β-catenin at the apico-lateral junction.

We predicted that phosphorylation of MCC by CK1δ/ε decreases the affinity of MCC to interacting partners at the centrosome, hence driving its relocalization to the apical ncMTOC in intestinal cells. To test this hypothesis, we overexpressed Myc-MCC singly or in combination with FLAG-CK1ε in HEK293 cells and immunoblotted for the centrosomal protein NDE1 along with the GTPase regulator protein RASAL2. We observed that in cells with CK1ε-induced MCC phosphorylation (+ Myc-MCC and + FLAG-CK1ε), the interaction with RASAL2 is stabilized, while binding to NDE1 is diminished, indicating a regulation of interactions between MCC and centrosomal proteins upon phosphorylation (Fig. *4D*).

We next generated small intestinal *ex vivo* organoids (Sato et al., 2009) and treated them with 5 μM of the potent CK1δ/ε-inhibitor (PF670462) on days 3, 4 and 5 post seeding (Fig. *4E,F*). Mcc localization was analyzed by IF in sections of inhibitor treated and non-treated control (NTC) organoids. Significantly, while Mcc signal was expectedly detected along the apical membrane of NTC organoids (Fig. *4G*), Mcc failed to relocalize to the ncMTOC upon inhibition of CK1δ/ε phosphorylation (Fig. *4H*). These results provide compelling evidence supporting our hypothesis that MCC is a direct target of CK1ε phosphorylation, which decreases the binding of MCC to centrosomal proteins such as NDE1 and triggers the relocalization of MCC to the apical membrane of differentiated intestinal cells (Fig. *4J, I*).

## Discussion

In this study, we extend our prior findings by comprehensively characterizing *Mcc* transcript expression and Mcc protein localization at the cellular level in the adult mouse small intestine (SI) and colon (Young et al., 2011). Our findings show that *Mcc* is broadly expressed in proliferating cells within intestinal crypts, extending distally from the CBC stem cells at the crypt base into the transit amplifying compartment. Notably, no expression of *Mcc* mRNA was observed in SI villi. Earlier studies using immunohistochemistry and immunoelectron microscopy described the presence of Mcc at the lateral membrane of murine SI epithelial cells and at the apical cytoplasm of colonic epithelial cells (Senda et al., 1999, 1997; Matsumine et al., 1996). Our IF and confocal microscopy analysis provide incontrovertible evidence that Mcc specifically localizes to the centrosome (MTOC) in crypt cells within the mouse and human intestinal epithelium. In contrast to *Mcc* mRNA distribution, the Mcc protein is additionally found in differentiated cells in the villus, explicitly localizing along the apical membrane—to the ncMTOC.

During intestinal cell differentiation, MTOC function is reassigned to the apical ncMTOC and is accompanied by the transcriptional downregulation of centrosomal genes and redeployment of centrosomal proteins to the apical cell membrane (Sen et al., 2010; Ito and Bettencourt-Dias, 2018; Muroyama and Lechler, 2017; Muroyama et al., 2018; Sanchez and Feldman, 2017). The relocalization of Mcc follows the intracellular trajectory of cardinal centrosomal proteins such as Ninein in intestinal cells (Goldspink et al., 2017b; Mogensen et al., 2000), and Pericentrin, which colocalizes with Mcc at the centrosome in the crypt and at the ncMTOC in the villus as shown in Fig. *3F* and Supp. Fig. *4F*. The shuttling of Ninein is proposed to occur via CLIP-170 to ncMTOCs, where it is captured by IQGAP1, an MCC-interacting protein (Fig. 2A) (Goldspink *et al*, 2017b). To date, however, the molecular mechanisms orchestrating redeployment of centrosomal proteins remain poorly characterized (Muroyama and Lechler, 2017; Sanchez and Feldman, 2017; Paz and Lüders, 2018; Gillard et al., 2021).

MCC is known a phosphoprotein, containing multiple potential serine and tyrosine phosphorylation sites (Pangon et al., 2012; Caria et al., 2019; Arnaud et al., 2009). In agreement with a large-scale protein–protein interaction screen in human cells, our proteomics studies confirm that MCC interacts with the serine/threonine kinases CK1δ/ε in HEK293 cells (Fig. 2A) (Ewing et al., 2007). Given the well-established role of casein kinases as intracellular effectors of the WNT signaling pathway (Gao et al., 2002; Cruciat, 2014; Su et al., 2018), we hypothesized that CK1 activation downstream of an unidentified WNT signal (“X” in Fig. *4I*) results in the phosphorylation of Mcc, triggering its relocalization to the ncMTOC during intestinal cellular differentiation. In support of this hypothesis, we first provide *in vitro* evidence that CK1ε phosphorylates MCC (Fig. *4A-C*). Second, in small intestinal organoids treated with the potent and specific CK1δ/ε inhibitor (PF670462) (Fig. *4G, H*), Mcc is no longer found apically in differentiated cells, but is dispersed throughout the cytoplasm. The phenotype observed in treated organoids (Fig. *4G*) can be attributed to the inhibition of Ck1δ/ε activity, as complete genetic loss of *Ck1δ/ε* in mice results in intestinal stem cell elimination, epithelial breakdown and rapid death (Morgenstern et al., 2017). These *in vivo* results guided both the low concentration of PF670462 (5 μM) used and the treatment window of our assay (Fig. 4EH). Future analyses are warranted to investigate the signaling network by which CK1δ/ε activity triggers the release-shuttle-capture of centrosomal proteins like MCC to the ncMTOC in differentiated intestinal cells.

Studies are additionally required to better establish the interactome of MCC in intestinal epithelial cells. One limitation of our proteomic studies is the use of the heterologous aneuploid HEK293 cell line, which is likely of adrenal origin (Lin et al., 2014). For example, such additional studies are necessary to identify endogenous intestinal MCC-binding proteins not recovered in HEK293 cells. One such protein is EZRIN, whose interaction with MCC and NHERF1 was previously reported (Reczek et al., 1997; Morales et al., 2004, 2007; Arnaud et al., 2009; Ewing et al., 2007). NHERF1 is a PDZ scaffold for the EZRIN-RADIXIN-MOESIN (ERM) protein family in epithelial cells (Solinet et al., 2013; Reczek et al., 1997). EZRIN is the only ERM found in the intestinal epithelium (Berryman et al., 1993; Reczek et al., 1997). The binding of EZRIN to NHERF1 via PDZ domains in intestinal cells is essential for the assembly of apical protein complexes, their interaction with the cytoskeleton and the establishment of cell polarity (Bretscher et al., 2002; Berryman et al., 1993; Garbett et al., 2010). β-catenin and YAP are known interactors of NHERF1 (Shibata et al., 2003; Mohler et al., 1999). Thus, we postulate that the PBM of MCC interacts with NHERF1-EZRIN at the apical membrane of intestinal cells and is potentially involved in establishing cell polarity and intracellular architecture (Fig 4G).

Homozygous *Mcc* null mutant mice were reported by us and others to be viable and fertile with no ostensible phenotypes (Young et al., 2011; Currey et al., 2019). However, the interactions of MCC and its cellular localization invite a more rigorous analysis of the intestinal epithelium in *Mcc*-deficient animals. For example, both *Nherf1*- and *Ezrin*-deficient mice present intestinal phenotypes, including polarity defects and epithelial cellular disorganization (Casaletto et al., 2011; Garbett et al., 2010; Morales et al., 2004; Saotome et al., 2004). Like *Mcc*, homozygous loss of *Nherf1* does not trigger adenoma formation along the length of the intestine (Young et al., 2011; Currey et al., 2019; Georgescu et al., 2016). However, in a recent study using the sulindac injury model, *Mcc* deficiency was shown to drive inflammation-associated colon cancer (Currey et al., 2019). Moreover, loss of *Nherf1* in combination with *Apc^Min/+^* resulted in a significant increase in tumor burden, followed by high levels of cytoplasmic β-catenin and nuclear YAP and upregulation of the WNT target gene *CyclinD1* (Georgescu et al., 2016). Given the prominent protein-protein interaction between NHERF1 and MCC and the tight genetic linkage between *APC* and *MCC* (Luongo et al., 1993), which is evolutionarily conserved suggesting that these two genes form a synexpression group (Niehrs and Pollet, 1999), it would be interesting to introduce the *Mcc* null mutation onto the sensitized, cancer-prone *Apc^Min^* genetic background (Luongo et al., 1993; Su et al., 1992). The generation of such mice may reveal subtle, hitherto unappreciated contributions of *Mcc* to intestinal tumor development.

## Materials and Methods

### Human tissues

Paraformaldehyde-Fixed Paraffin-Embedded (PFPE) human intestinal tissue sections were provided by Assoc. Prof. Christopher S. Williams. Tissues were acquired following all ethical regulations of the Department of Medicine and Cancer Biology at Vanderbilt University School of Medicine - Nashville, TN - USA.

### Animals

Mice used for this study were housed, bred and euthanised according to the Institutional Animal Care and Use Committee (IACUC) of Singapore under the protocol number #A20027. This study was performed following all ethical regulations of the Animal Research Facility (ARF) from the Lee Kong Chian School of Medicine (LKCMedicine), Nanyang Technological University (NTU). Experiments were conducted using animals with a minimum age of 90 days. *Mcc^lacZ^* mutant mice were genotyped as previously described (Young et al., 2011).

### Mouse small intestine villi and crypts isolation

Villus and crypt compartments from mouse duodenum were dissociated as previously described (Sato et al., 2009; Goldspink et al., 2017). Briefly, duodenum (8 - 10 cm) segments were washed with ice cold 1X PBS-0 (lacking Mg^2+^ and Ca^2+^). Specimens were cut open longitudinally. Villus fractions were harvested and collected into 50 ml Falcon conical tubes with ice-cold PBS-0. Crypt fractions were filtered through a 70 μm cell strainer (Biosciences #352350). Purified crypt and villus fractions were further processed either for RNA isolation, protein extraction, immunohistochemistry or 3D organoid culture.

### Intestinal organoid culture

About 50 isolated crypts/well were seeded in 50 μl of Matrigel (Corning #356231) and cultured in 48-well plates (Corning cat# 3526) using Mouse IntestiCult™ Organoid Growth Medium (Stemcell Technologies #06005) supplemented with 100 mg/ml ampicillin (Sigma #A0166) and 100mg/mL Primocin (Invivogen #ANTPM1) following manufacturer’s instructions. Only primary cells from mice were used for organoid culture. For inhibition of phosphorylation, a single dose (5 μM) of the Casein Kinase 1 δ/ε inhibitor PF670462 (Sigma #SML0795) was added to the medium on days 3, 4 and 5 of culture post seeding. Treated and non-treated control organoids were harvested on day 6 for histological analysis.

### Cell culture

Retinal pigment epithelial (RPE-1) and human embryonic kidney (HEK-293) cells were cultured in Dulbecco’s Modified Eagle Medium (DMEM) High glucose, (Gibco #11960-044) supplemented with 10% (v/v) Fetal Bovine Serum (FBS) (Sigma #F2442). Human colon carcinoma HCT116 cells were cultured in McCoy’s 5A Modified Medium (Gibco #16600-082) supplemented with 10% (v/v) FBS. Human colon cancer SW480 cells were cultured in RPMI medium supplemented with 10% (v/v) FBS, 2 mM L-glutamine (Gibco #25030-081), 1 mM sodium pyruvate (Gibco #11360-070) and Penicillin-Streptomycin (Gibco #15140-122).

### Histology

The proximal portion of the small intestine (duodenum) and distal portion of the colon were used for the histological analyses. Tissues were carefully flushed with ice-cold PBS, cut open longitudinally, and fixed overnight at 4°C with 4% Paraformaldehyde (PFA) (EMS #15710). Specimens (roughly 3-cm pieces) were later dehydrated and embedded in paraffin. For wholemount tissue analysis, samples were fixed overnight with 4% PFA and washed with PBS. Staining for β-galactosidase activity was performed as described in Merkwitz *et al* (2016). Alkaline Phosphatase staining was performed using the Vulcan Fast Red Chromogen kit (Biocare Medical #FR805S). ISH was performed using RNAscope®2.5 HD Reagent Kit-Brown (#322300) and 2.5 HD Duplex Reagent Kit (#322430) from Advanced Cell Diagnostics (ACD). The specificity of the Mm*Mcc* probe set (ACD #411961 and #411961-C2) was confirmed by lack of signal in sections of *Mcc^lacZ^* homozygous adult intestine. Other probes: *Mm-Lgr5* (ACD #312171 and #312171-C2) and Mm-*Ppib* (ACD #313911). For immunofluorescence, tissue sections (7 μm) were deparaffinized and rehydrated. Antigen retrieval was performed with 0.5x Tris/Borate/EDTA buffer using a pressure cooker. Sections were then blocked with 5% Bovine Serum Albumin (BSA) (Sigma #A3311) for 2 hours at room temperature (RT) and stained overnight with primary antibodies (Table 01). Tissues were washed with PBS before incubation with secondary antibodies for 1 hour at RT. Nuclei were stained with Hoechst 33342 (Thermo Fisher #62249). Slides were mounted with water-based mounting medium Hydromount (EMS #17966). Representative images of 3 repeats were included in the manuscript.

### Immunofluorescence of cells

RPE-1, HCT116, HEK293 and SW480 cells were grown on glass coverslips and fixed in cold 100% methanol for 20 mins at −20°C. Cells were washed/permeabilized with 0.5% Triton-X100 (Sigma #X100) in PBS and then blocked for 30 minutes with 5% BSA in 0.1% Triton-X100 in PBS. Immunostaining with primary antibodies (Table 01) was performed over night at RT followed by PBS washes. Incubation with secondary antibody (Table 02) was performed for 2 hours at room temperature. Nuclei were stained using Hoechst 33342. Slides were mounted with water-based mounting medium Hydromount. For Structured illumination microscopy (SIM), cells were grown in 1.5H circular glass coverslip, stained, and mounted with VECTASHIELD mounting medium (Vector Laboratories #H-1000).

**Table 02.** Secondary Antibodies

| Antibody | Species | Catalog # | Brand | Application |
| --- | --- | --- | --- | --- |
| Alexa Fluor 488 Anti-Mouse | Donkey | A21202 | Invitrogen | IF |
| Alexa Fluor 488 Anti-Rabbit | Donkey | A21206 | Invitrogen | IF |
| Alexa Fluor 594 Anti-Mouse | Donkey | A21203 | Invitrogen | IF |
| Alexa Fluor 594 Anti-Rabbit | Donkey | A21207 | Invitrogen | IF |
| Alexa Fluor 488 Anti-Mouse | Goat | A28175 | Invitrogen | IF |

### Imaging

Confocal microscopy images were acquired either using Olympus FV1000 upright, Olympus FV3000 laser scanning, or Zeiss LSM800 Airyscan microscopes. DeltaVision OMX 3D-SIM Oxford Nanoimager microscope was used for SIM imaging. Conventional bright-field images were taken using Zeiss AxioImager Z1 upright (ZAZ1). Images were further processed using FiJi (ImageJ). The instruments used are either from the A*STAR Microscopy Platform (FV1000, FV3000, ZAZ1 and OMX) or from the LKCMedicine – NTU (LSM800).

### RNA isolation and quantitative PCR (qPCR)

RNA extraction from tissues was performed using TRIzol (Invitrogen #15596026) and RNeasy kit (Qiagen # 74004). High-Capacity cDNA Reverse Transcription Kit (Applied Biosystems #4368814) was used for cDNA synthesis. Quantification of gene expression by q-PCR was performed using Power SYBR Green PCR Master Mix (Applied Biosystems #4309155). Technical triplicates for a minimum of three biological replicates were used. Relative gene expression was assessed using double CT method and data were normalized with *β-actin*. Data were tested for significance by an unpaired two-tailed t-test. P-values of statistical significance are represented as ***p < 0.001, **p < 0.01. The qPCR primers are listed in Table 03.

**Table 03.**
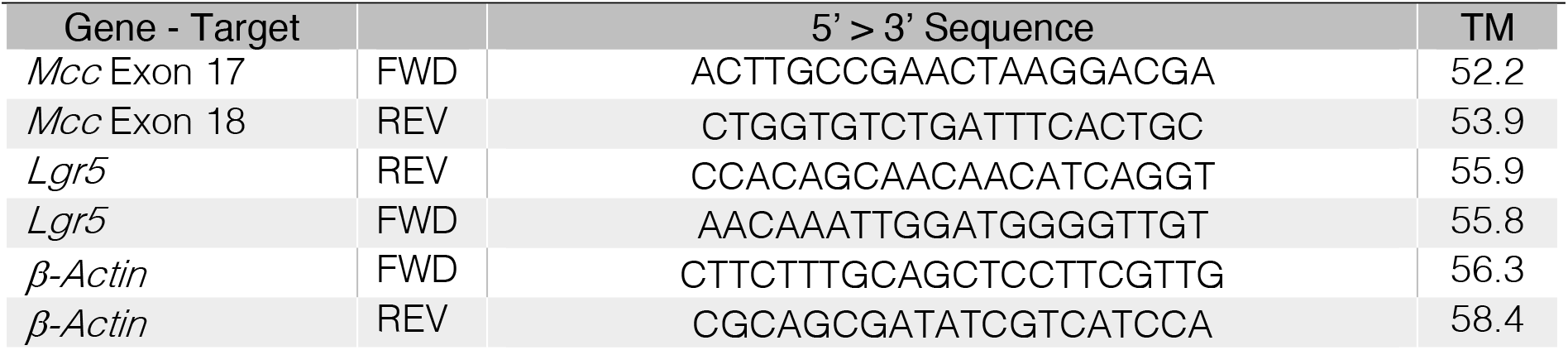
qPCR Primers.

### Protein extraction and Western Blot

Tissue/cell lysates were prepared using RIPA (SIGMA #R0278) lysis buffer supplemented with cOmplete-ULTRA protease inhibitor (Roche #05892970001). Tissue lysates were later sonicated for two cycles at 4°C for complete lysis. Protein concentration was determined using the Bradford assay. Samples were boiled at 95°C for 5 min. After electrophoresis, proteins were transferred onto a PVDF membrane (Millipore #ISEQ20200) and incubated 1 hour at room temperature with 2% milk powder in PBS, 0.2% TRITON 100 for blocking. Primary antibodies were applied overnight at 4°C and secondary antibodies for 1 hour at room temperature. Blots were revealed using SuperSignal (Thermo Scientific #34095). β-actin was routinely used as a loading control.

### Constructs

Full length cDNA constructs of human MCC (P23508-1), NDE1 and RASAL2 were PCR-cloned into pCMV2B (Flag) and pCMV3B (Myc) constructs (Stratagene). Expression constructs for SCRIBBLE, NHERF1, and CK1E, were generated by Gateway Cloning into pcDNA5 FRT-TO (3X FLAG N-terminus or EGFP N-terminus) vectors (Invitrogen). Point mutations and truncations were generated by site-directed mutagenesis using the QuikChange II Site-Directed mutagenesis kit (Agilent #200524) and verified by sequencing. The pCS2+_EmGFP-Mcc (G3UW40) was previously described in Young et al., 2014. This plasmid robustly produces an N-terminal EmGFP-Mcc fusion protein in heterologous cells. (Plasmid inventory #525, Dunn lab). HEK293 cells were transfected with a minimum amount of EmGFP-Mcc yielding a mosaic EmGFP-Mcc signal in Figure *2C*.

### Immunoprecipitation and Mass-spectrometry

For immunoprecipitation, cells were lysed in buffer containing 50 mM HEPES (pH 7.5), 150 mM NaCl, 1% Triton X-100, 10% glycerol, and 1 mM EDTA. Lysates, precleared with protein G-Sepharose 4 beads (GE Healthcare #17-0618-0), were incubated with an anti-Flag monoclonal antibody or anti-GFP polyclonal for 1 hr, followed by incubation with protein G-Sepharose beads for 1 hr. The beads were subsequently washed three times in the lysis buffer. MCC interaction partners were captured using two approaches (Gel-free and Gel-digest). In both, N-terminal FLAG-tagged human MCC (P23508-1) constructs (pCMV2B vector) were transfected into HEK293 cells and selected with G418 (Gibco #10131035). G418-resistant cells were grown, lysed and immunoprecipitated using anti-FLAG agarose beads. Cells stably expressing an empty vector were used as control. To capture interactions, samples were run either on an SDS-Page gel (Gel-digested approach) or eluted using 200 mM glycine pH 2.5 (Gel-free approach). Samples eluted with glycine were subjected to separation with SCX (strong-cation exchange) column (Sepax Tech #Z777148) followed by digestion with trypsin (Gibco #R001100). SDS-Page gel samples were run against an empty vector control (*pCMV* empty) and bands present in the FLAG-MCC lane were extracted and digested. Samples from both approaches were subjected to mass spectrometry (OrbiTrap Elite) and analysed using Mascot (Matrix Science) and ProHITS (Samuel Lunenfeld Research Institute, Mount Sinai Hospital, ON, Canada) for protein identification and quantification. The Human NCBI Reference Sequence (RefSeq NCBI) database was searched by Mascot with the following parameters: mass tolerance: 7 ppm on MS and 0.5 Daltons on MS/MS; maximum missed cleavages: 2; fixed modification: carbamidomethyl (C); variable modification:oxidation (M), phospho (ST) and phospho (Y). ProHITS was utilized to filter out common contaminant interactions from FLAG-tag affinity based on cRAP (common Repository of Adventitious Proteins) database.

## Acknowledgments

We thank Drs. Radoslaw Sobota (IMCB, Singapore) and Bernett Lee (SIgN, Singapore) for their invaluable assistance with bioinformatics analyses, Drs. Brian Burke and Lee Yin Loon (SRIS, Singapore) for insightful discussions about centrosome biology, Dr. Graham Wright (A*STAR Microscopy Platform) for technical assistance with SIM, Drs. Nicholas Barker (IMCB, Singapore) and Kazuhiro Murakami (Kanazawa University, Japan) for both their helpful discussions and reagents supporting intestinal organoid culture and analysis, and Dr. Brigid Hogan for comments on the manuscript. This work was funded by the Institute of Medical Biology (A*STAR, Singapore) as well as start-up funding provided by the Lee Kong Chian School of Medicine, Nanyang Technological University and the Ministry of Education (MOE), Singapore (Continuation Grant—Endodermal Development and Differentiation (EDD) Lab) to N.R.D.. L.B.T. was initially funded by the Singapore International Graduate Award (SINGA), A*STAR Graduate Academy (A*GA). B.A.L. was funded by the Canadian Institute of Health Research Postdoctoral Fellowship. Lastly, we wish to honour the legacy and significant scientific contributions of the late Professor Tony Pawson (RIP 2013) in whose lab B.A.L. first discovered the centrosomal localization of MCC.

## Author Contributions

N.R.D., B.A.L. and L.B.T. designed research; L.B.T. and B.A.L., performed research; N.R.D., L.B.T., B.A.L., A.-C.G., and M.L. analysed data; S.L.M.O., E.K.T., M.M., C.Y., N.S.T., C.S.W., M.L. contributed to the research; L.B.T., M.L., N.R.D. wrote the paper.

## Classification

Biological Sciences; Cell Biology

## References

Arnaud, C., M. Sebbagh, S. Nola, S. Audebert, G. Bidaut, A. Hermant, O. Gayet, N.J. Dusetti, V. Ollendorff, M.-J. Santoni, J.-P. Borg, and P. Lécine. 2009. MCC, a new interacting protein for Scrib, is required for cell migration in epithelial cells. Febs Lett. 583:2326–32. doi:10.1016/j.febslet.2009.06.034.

Badura, L., T. Swanson, W. Adamowicz, J. Adams, J. Cianfrogna, K. Fisher, J. Holland, R. Kleiman, F. Nelson, L. Reynolds, K.S. Germain, E. Schaeffer, B. Tate, and J. Sprouse. 2007. An Inhibitor of Casein Kinase Iε Induces Phase Delays in Circadian Rhythms under Free-Running and Entrained Conditions. J Pharmacol Exp Ther. 322:730–738. doi:10.1124/jpet.107.122846.

Barker, N. 2014. Adult intestinal stem cells: critical drivers of epithelial homeostasis and regeneration. Nat Rev Mol Cell Bio. 15:19–33. doi:10.1038/nrm3721.

Barker, N., J.H. van Es, J. Kuipers, P. Kujala, M. van den Born, M. Cozijnsen, A. Haegebarth, J. Korving, H. Begthel, P.J. Peters, and H. Clevers. 2007. Identification of stem cells in small intestine and colon by marker gene Lgr5. Nature. 449:1003–1007. doi:10.1038/nature06196.

Benthani, F.A., D. Herrmann, P.N. Tran, L. Pangon, M.C. Lucas, A.H. Allam, N. Currey, S. Al-Sohaily, M. Giry-Laterriere, J. Warusavitarne, P. Timpson, and M.R.J. Kohonen-Corish. 2018. ‘MCC’ protein interacts with E-cadherin and β-catenin strengthening cell–cell adhesion of HCT116 colon cancer cells. Oncogene. 37:663–672. doi:10.1038/onc.2017.362.

Berryman, M., Z. Franck, and A. Bretscher. 1993. Ezrin is concentrated in the apical microvilli of a wide variety of epithelial cells whereas moesin is found primarily in endothelial cells. J Cell Sci. 105 ( Pt 4):1025–43.

Bouckson-Castaing, V., M. Moudjou, D.J. Ferguson, S. Mucklow, Y. Belkaid, G. Milon, and P.R. Crocker. 1996. Molecular characterisation of ninein, a new coiled-coil protein of the centrosome. J Cell Sci. 109:179–190. doi:10.1242/jcs.109.1.179.

Bourne, H.R. 1991. Consider the coiled coil. Nature. 351:188–189. doi:10.1038/351188a0.

Bretscher, A., K. Edwards, and R.G. Fehon. 2002. ERM proteins and merlin: integrators at the cell cortex. Nat Rev Mol Cell Bio. 3:586–599. doi:10.1038/nrm882.

Brinkley, B.R. 1985. Microtubule Organizing Centers. Annu Rev Cell Dev Bi. 1:145–172. doi:10.1146/annurev.cb.01.110185.001045.

Caria, S., B.Z. Stewart, R. Jin, B.J. Smith, P.O. Humbert, and M. Kvansakul. 2019. Structural analysis of phosphorylation-associated interactions of human MCC with Scribble PDZ domains. Febs J. doi:10.1111/febs.15002.

Casaletto, J.B., I. Saotome, M. Curto, and A.I. McClatchey. 2011. Ezrin-mediated apical integrity is required for intestinal homeostasis. Proc National Acad Sci. 108:11924–11929. doi:10.1073/pnas.1103418108.

Clevers, H. 2013. The Intestinal Crypt, A Prototype Stem Cell Compartment. Cell. 154:274–284. doi:10.1016/j.cell.2013.07.004.

Cruciat, C.-M. 2014. Casein kinase 1 and Wnt/β-catenin signaling. Curr Opin Cell Biol. 31:46–55. doi:10.1016/j.ceb.2014.08.003.

Currey, N., Z. Jahan, C.E. Caldon, P.N. Tran, F. Benthani, P.D. Lacavalerie, D.L. Roden, B.S. Gloss, C. Campos, E.G. Bean, A. Bullman, S. Reibe-Pal, M.E. Dinger, M.A. Febbraio, S.J. Clarke, J.E. Dahlstrom, and M.R.J. Kohonen-Corish. 2019. Mouse model of “Mutated in Colorectal Cancer” gene deletion reveals novel pathways in inflammation and cancer. Cell Mol Gastroenterology Hepatology. 7:819–839. doi:10.1016/j.jcmgh.2019.01.009.

Doxsey, S.J., P. Stein, L. Evans, P.D. Calarco, and M. Kirschner. 1994. Pericentrin, a highly conserved centrosome protein involved in microtubule organization. Cell. 76:639–650. doi:10.1016/0092-8674(94)90504-5.

Ewing, R.M., P. Chu, F. Elisma, H. Li, P. Taylor, S. Climie, L. McBroom-Cerajewski, M.D. Robinson, L. O’Connor, M. Li, R. Taylor, M. Dharsee, Y. Ho, A. Heilbut, L. Moore, S. Zhang, O. Ornatsky, Y.V. Bukhman, M. Ethier, Y. Sheng, J. Vasilescu, M. Abu-Farha, J. Lambert, H.S. Duewel, I.I. Stewart, B. Kuehl, K. Hogue, K. Colwill, K. Gladwish, B. Muskat, R. Kinach, S. Adams, M.F. Moran, G.B. Morin, T. Topaloglou, and D. Figeys. 2007. Large-scale mapping of human protein–protein interactions by mass spectrometry. Mol Syst Biol. 3:89. doi:10.1038/msb4100134.

Fukuyama, R., R. Niculaita, K.P. Ng, E. Obusez, J. Sanchez, M. Kalady, P.P. Aung, G. Casey, and N. Sizemore. 2008. Mutated in colorectal cancer, a putative tumor suppressor for serrated colorectal cancer, selectively represses β-catenin-dependent transcription. Oncogene. 27:6044–6055. doi:10.1038/onc.2008.204.

Gao, Z.-H., J.M. Seeling, V. Hill, A. Yochum, and D.M. Virshup. 2002. Casein kinase I phosphorylates and destabilizes the β-catenin degradation complex. Proc National Acad Sci. 99:1182–1187. doi:10.1073/pnas.032468199.

Garbett, D., D.P. LaLonde, and A. Bretscher. 2010. The scaffolding protein EBP50 regulates microvillar assembly in a phosphorylation-dependent manner. J Cell Biology. 191:397–413. doi:10.1083/jcb.201004115.

Gehart, H., and H. Clevers. 2019. Tales from the crypt: new insights into intestinal stem cells. Nat Rev Gastroentero. 16:19–34. doi:10.1038/s41575-018-0081-y.

Georgescu, M.-M., M. Gagea, and G. Cote. 2016. NHERF1/EBP50 Suppresses Wnt-β-Catenin Pathway–Driven Intestinal Neoplasia. Neoplasia. 18:512–523. doi:10.1016/j.neo.2016.07.003.

Gillard, G., G. Girdler, and K. Röper. 2021. A release-and-capture mechanism generates an essential non-centrosomal microtubule array during tube budding. Nat Commun. 12:4096. doi:10.1038/s41467-021-24332-0.

Goldspink, D.A., Z.J. Matthews, E.K. Lund, T. Wileman, and M.M. Mogensen. 2017a. Immuno-fluorescent Labeling of Microtubules and Centrosomal Proteins in <em>Ex Vivo</em> Intestinal Tissue and 3D <em>In Vitro</em> Intestinal Organoids. J Vis Exp.doi:10.3791/56662.

Goldspink, D.A., C. Rookyard, B.J. Tyrrell, J. Gadsby, J. Perkins, E.K. Lund, N. Galjart, P. Thomas, T. Wileman, and M.M. Mogensen. 2017b. Ninein is essential for apico-basal microtubule formation and CLIP-170 facilitates its redeployment to non-centrosomal microtubule organizing centres. Open Biol. 7:160274. doi:10.1098/rsob.160274.

Greer, Y.E., C.J. Westlake, B. Gao, K. Bharti, Y. Shiba, C.P. Xavier, G.J. Pazour, Y. Yang, and J.S. Rubin. 2014. Casein kinase 1δ functions at the centrosome and Golgi to promote ciliogenesis. Mol Biol Cell. 25:1629–1640. doi:10.1091/mbc.e13-10-0598.

Groden, J., A. Thliveris, W. Samowitz, M. Carlson, L. Gelbert, H. Albertsen, G. Joslyn, J. Stevens, L. Spirio, M. Robertson, L. Sargeant, K. Krapcho, E. Wolff, R. Burt, J.P. Hughes, J. Warrington, J. McPherson, J. Wasmuth, D.L. Paslier, H. Abderrahim, D. Cohen, M. Leppert, and R. White. 1991. Identification and characterization of the familial adenomatous polyposis coli gene. Cell. 66:589–600. doi:10.1016/0092-8674(81)90021-0.

Heintzmann, R., and T. Huser. 2017. Super-Resolution Structured Illumination Microscopy. Chem Rev. 117:13890–13908. doi:10.1021/acs.chemrev.7b00218.

Hinnebusch, B.F., A. Siddique, J.W. Henderson, M.S. Malo, W. Zhang, C.P. Athaide, M.A. Abedrapo, X. Chen, V.W. Yang, and R.A. Hodin. 2004. Enterocyte differentiation marker intestinal alkaline phosphatase is a target gene of the gut-enriched Krüppel-like factor. Am J Physiol-gastr L. 286:G23–G30. doi:10.1152/ajpgi.00203.2003.

Hua, K., and R.J. Ferland. 2017. Fixation methods can differentially affect ciliary protein immunolabeling. Cilia. 6:5. doi:10.1186/s13630-017-0045-9.

Ito, D., and M. Bettencourt-Dias. 2018. Centrosome Remodelling in Evolution. Cells. 7:71. doi:10.3390/cells7070071.

Keenan, C.R., S.Y. Langenbach, F. Jativa, T. Harris, M. Li, Q. Chen, Y. Xia, B. Gao, M.J. Schuliga, J. Jaffar, D. Prodanovic, Y. Tu, A. Berhan, P.V.S. Lee, G.P. Westall, and A.G. Stewart. 2018. Casein Kinase 1δ/ε Inhibitor, PF670462 Attenuates the Fibrogenic Effects of Transforming Growth Factor-β in Pulmonary Fibrosis. Front Pharmacol. 9:738. doi:10.3389/fphar.2018.00738.

Kinzler, K., M. Nilbert, L. Su, B. Vogelstein, T. Bryan, D. Levy, K. Smith, A. Preisinger, P. Hedge, D. McKechnie, and al. et. 1991a. Identification of FAP locus genes from chromosome 5q21. Science. 253:661–665. doi:10.1126/science.1651562.

Kinzler, K., M. Nilbert, B. Vogelstein, T. Bryan, D. Levy, K. Smith, A. Preisinger, Hamilton, P. Hedge, A. Markham, and al. et. 1991b. Identification of a gene located at chromosome 5q21 that is mutated in colorectal cancers. Science. 251:1366–1370. doi:10.1126/science.1848370.

Leushacke, M., and N. Barker. 2014. Ex vivo culture of the intestinal epithelium: strategies and applications. Gut. 63:1345. doi:10.1136/gutjnl-2014-307204.

Lin, Y.-C., M. Boone, L. Meuris, I. Lemmens, N.V. Roy, A. Soete, J. Reumers, M. Moisse, S. Plaisance, R. Drmanac, J. Chen, F. Speleman, D. Lambrechts, Y.V. de Peer, J. Tavernier, and N. Callewaert. 2014. Genome dynamics of the human embryonic kidney 293 lineage in response to cell biology manipulations. Nat Commun. 5:4767. doi:10.1038/ncomms5767.

Luongo, C., K.A. Gould, L.-K. Su, K.W. Kinzler, B. Vogelstein, W. Dietrich, E.S. Lander, and A.R. Moser. 1993. Mapping of Multiple Intestinal Neoplasia (Min) to Proximal Chromosome 18 of the Mouse. Genomics. 15:3–8. doi:10.1006/geno.1993.1002.

Matsumine, A., T. Senda, G.-H. Baeg, B.C. Roy, Y. Nakamura, M. Noda, K. Toyoshima, and T. Akiyama. 1996. MCC, a Cytoplasmic Protein That Blocks Cell Cycle Progression from the G/G to S Phase. J Biol Chem. 271:10341–10346. doi:10.1074/jbc.271.17.10341.

Meads, T., and T.A. Schroer. 1995. Polarity and nucleation of microtubules in polarized epithelial cells. Cell Motil Cytoskel. 32:273–288. doi:10.1002/cm.970320404.

Merkwitz, C., O. Blaschuk, A. Schulz, and A.M. Ricken. 2016. Comments on Methods to Suppress Endogenous β-Galactosidase Activity in Mouse Tissues Expressing the LacZ Reporter Gene. J Histochem Cytochem. 64:579–586. doi:10.1369/0022155416665337.

Mogensen, M.M., A. Malik, M. Piel, V. Bouckson-Castaing, and M. Bornens. 2000. Microtubule minus-end anchorage at centrosomal and non-centrosomal sites: the role of ninein. J Cell Sci. 113 ( Pt 17):3013–23.

Mohler, P.J., S.M. Kreda, R.C. Boucher, M. Sudol, M.J. Stutts, and S.L. Milgram. 1999. Yes-Associated Protein 65 Localizes P62c-Yes to the Apical Compartment of Airway Epithelia by Association with Ebp50. J Cell Biology. 147:879–890. doi:10.1083/jcb.147.4.879.

Morales, F.C., Y. Takahashi, E.L. Kreimann, and M.-M. Georgescu. 2004. Ezrin-radixin-moesin (ERM)-binding phosphoprotein 50 organizes ERM proteins at the apical membrane of polarized epithelia. Proc National Acad Sci. 101:17705–17710. doi:10.1073/pnas.0407974101.

Morales, F.C., Y. Takahashi, S. Momin, H. Adams, X. Chen, and M.-M. Georgescu. 2007. NHERF1/EBP50 Head-to-Tail Intramolecular Interaction Masks Association with PDZ Domain Ligands. Mol Cell Biol. 27:2527–2537. doi:10.1128/mcb.01372-06.

Morgenstern, Y., U.D. Adhikari, M. Ayyash, E. Elyada, B. Tóth, A. Moor, S. Itzkovitz, and Y. Ben-Neriah. 2017. Casein kinase 1-epsilon or 1-delta required for Wnt-mediated intestinal stem cell maintenance. Embo J. 36:3046–3061. doi:10.15252/embj.201696253.

Muroyama, A., and T. Lechler. 2017. Microtubule organization, dynamics and functions in differentiated cells. Development. 144:3012–3021. doi:10.1242/dev.153171.

Muroyama, A., M. Terwilliger, B. Dong, H. Suh, and T. Lechler. 2018. Genetically induced microtubule disruption in the mouse intestine impairs intracellular organization and transport. Mol Biol Cell. 29:1533–1541. doi:10.1091/mbc.e18-01-0057.

Niehrs, C., and N. Pollet. 1999. Synexpression groups in eukaryotes. Nature. 402:483–7. doi:10.1038/990025.

Nishisho, I., Y. Nakamura, Y. Miyoshi, Y. Miki, H. Ando, A. Horii, K. Koyama, J. Utsunomiya, S. Baba, and P. Hedge. 1991. Mutations of chromosome 5q21 genes in FAP and colorectal cancer patients. Science. 253:665–669. doi:10.1126/science.1651563.

Pangon, L., C.V. Kralingen, M. Abas, R.J. Daly, E.A. Musgrove, and M.R.J. Kohonen-Corish. 2012. The PDZ-binding motif of MCC is phosphorylated at position −1 and controls lamellipodia formation in colon epithelial cells. Biochimica Et Biophysica Acta Bba - Mol Cell Res. 1823:1058–1067. doi:10.1016/j.bbamcr.2012.03.011.

Pangon, L., D. Mladenova, L. Watkins, C. Kralingen, N. Currey, S. Al-Sohaily, P. Lecine, J. Borg, and M.R.J. Kohonen-Corish. 2015. MCC inhibits beta-catenin transcriptional activity by sequestering DBC1 in the cytoplasm. Int J Cancer. 136:55–64. doi:10.1002/ijc.28967.

Pangon, L., N.D. Sigglekow, M. Larance, S. Al-Sohaily, D.N. Mladenova, C.I. Selinger, E.A. Musgrove, and M.R.J. Kohonen-Corish. 2010. The “Mutated in Colorectal Cancer” Protein Is a Novel Target of the UV-Induced DNA Damage Checkpoint. Genes Cancer. 1:917–926. doi:10.1177/1947601910388937.

Paz, J., and J. Lüders. 2018. Microtubule-Organizing Centers: Towards a Minimal Parts List. Trends Cell Biol. 28:176–187. doi:10.1016/j.tcb.2017.10.005.

Peters, J.M., R.M. McKay, J.P. McKay, and J.M. Graff. 1999. Casein kinase I transduces Wnt signals. Nature. 401:345–350. doi:10.1038/43830.

Reczek, D., M. Berryman, and A. Bretscher. 1997. Identification of EBP50: A PDZ-containing Phosphoprotein that Associates with Members of the Ezrin-Radixin-Moesin Family. J Cell Biology. 139:169–179. doi:10.1083/jcb.139.1.169.

Sakanaka, C., P. Leong, L. Xu, S.D. Harrison, and L.T. Williams. 1999. Casein kinase Iε in the Wnt pathway: Regulation of β-catenin function. Proc National Acad Sci. 96:12548–12552. doi:10.1073/pnas.96.22.12548.

Sanchez, A.D., and J.L. Feldman. 2017. Microtubule-organizing centers: from the centrosome to non-centrosomal sites. Curr Opin Cell Biol. 44:93–101. doi:10.1016/j.ceb.2016.09.003.

Saotome, I., M. Curto, and A.I. McClatchey. 2004. Ezrin Is Essential for Epithelial Organization and Villus Morphogenesis in the Developing Intestine. Dev Cell. 6:855–864. doi:10.1016/j.devcel.2004.05.007.

Sato, T., R.G. Vries, H.J. Snippert, M. van de Wetering, N. Barker, D.E. Stange, J.H. van Es, A. Abo, P. Kujala, P.J. Peters, and H. Clevers. 2009. Single Lgr5 stem cells build crypt-villus structures in vitro without a mesenchymal niche. Nature. 459:262–265. doi:10.1038/nature07935.

Sen, G.L., J.A. Reuter, D.E. Webster, L. Zhu, and P.A. Khavari. 2010. DNMT1 maintains progenitor function in self-renewing somatic tissue. Nature. 463:563–567. doi:10.1038/nature08683.

Senda, T., A. Matsumine, T. Akiyama, and S. Kobayashi. 1997. Association of the MCC gene product with the plasma membrane and membrane organelles. Medical Electron Microsc. 30:1–7. doi:10.1007/bf01458345.

Senda, T., A. Matsumine, H. Yanai, and T. Akiyama. 1999. Localization of MCC (Mutated in Colorectal Cancer) in Various Tissues of Mice and Its Involvement in Cell Differentiation. J Histoohem Cytoohem. 47:1149–1157. doi:10.1177/002215549904700907.

Shibata, T., M. Chuma, A. Kokubu, M. Sakamoto, and S. Hirohashi. 2003. EBP50, a β-catenin-associating protein, enhances Wnt signaling and is over-expressed in hepatocellular carcinoma. Hepatology. 38:178–186. doi:10.1053/jhep.2003.50270.

Solinet, S., K. Mahmud, S.F. Stewman, K.B.E. Kadhi, B. Decelle, L. Talje, A. Ma, B.H. Kwok, and S. Carreno. 2013. The actin-binding ERM protein Moesin binds to and stabilizes microtubules at the cell cortex. J Cell Biology. 202:251–60. doi:10.1083/jcb.201304052.

Su, L., K. Kinzler, B. Vogelstein, A. Preisinger, A. Moser, C. Luongo, K. Gould, and W. Dove. 1992. Multiple intestinal neoplasia caused by a mutation in the murine homolog of the APC gene. Science. 256:668–670. doi:10.1126/science.1350108.

Su, Z., J. Song, Z. Wang, L. Zhou, Y. Xia, S. Yu, Q. Sun, S.-S. Liu, L. Zhao, S. Li, L. Wei, D.A. Carson, and D. Lu. 2018. Tumor promoter TPA activates Wnt/β-catenin signaling in a casein kinase 1-dependent manner. Proc National Acad Sci. 115:201802422. doi:10.1073/pnas.1802422115.

Sussman, N.L., R. Eliakim, D. Rubin, D.H. Perlmutter, K. DeSchryver-Kecskemeti, and D.H. Alpers. 1989. Intestinal alkaline phosphatase is secreted bidirectionally from villous enterocytes. Am J Physiol-gastr L. 257:G14–G23. doi:10.1152/ajpgi.1989.257.1.g14.

Waller, A., S. Findeis, and M. Lee. 2016. Familial Adenomatous Polyposis. J Pediatric Genetics. 05:078–083. doi:10.1055/s-0036-1579760.

Young, T., Y. Poobalan, Y. Ali, W.S. Tein, A. Sadasivam, T.E. Kim, P.E. Tay, and N.R. Dunn. 2011. Mutated in colorectal cancer (Mcc), a candidate tumor suppressor, is dynamically expressed during mouse embryogenesis. Dev Dynam. 240:2166–2174. doi:10.1002/dvdy.22712.

