## Supplementary Figures for "Mutated in Colorectal Cancer (MCC) is a centrosomal protein that relocalizes to the ncMTOC during intestinal cell differentiation"

#### Supp Figure 01

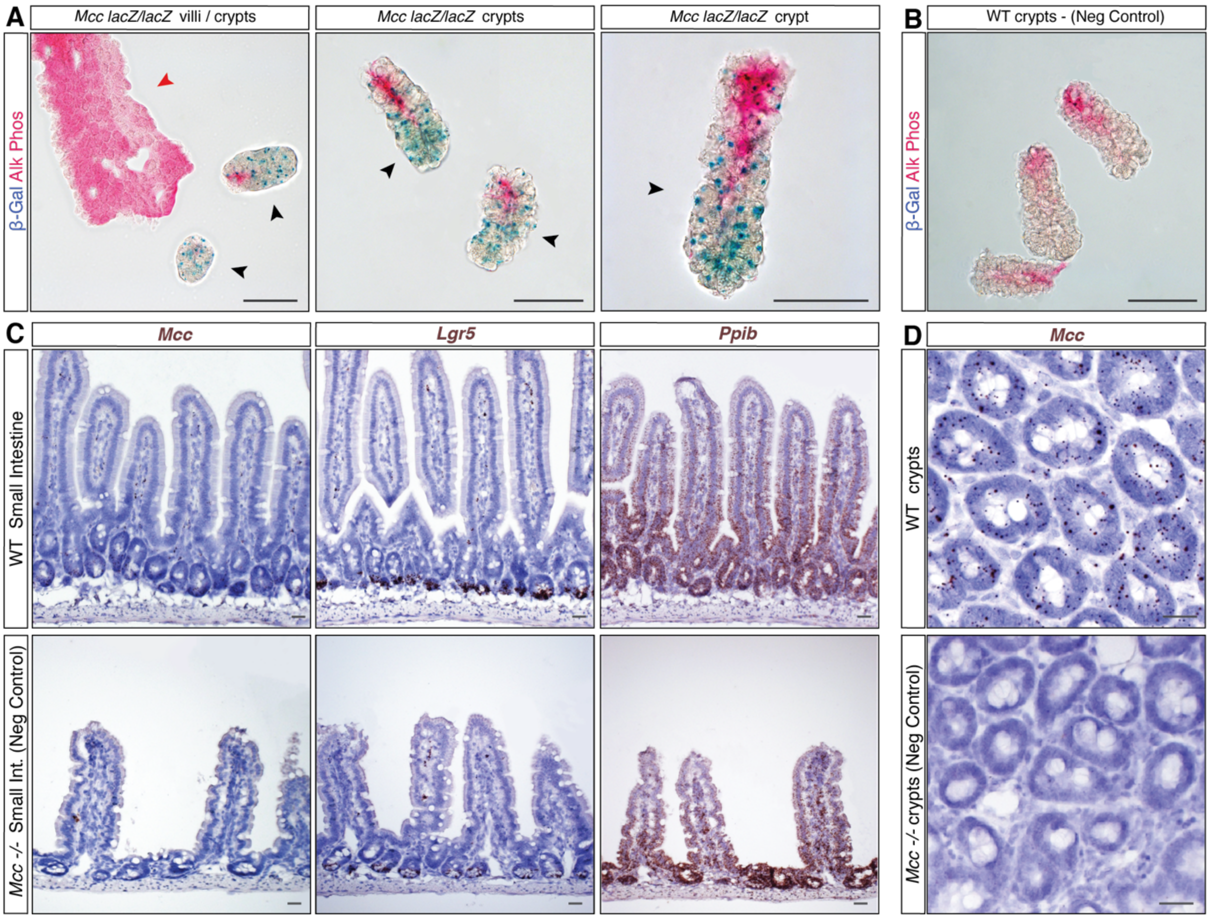


**Supplementary Figure 1. β-Galactosidase (β-Gal) staining activity and *in situ* hybridization (ISH) controls.** *(A):* Co-staining for β-Gal (blue) activity and intestinal alkaline phosphatase (red) in whole-mount villus (red arrowhead) and crypt (black arrowheads) fractions isolated from *Mcc*$^{lacZ/lacZ}$ adult small intestine (SI). *(B):* Wild-type (WT) β-Gal-negative crypt controls. Scale bars, 50 µm. *(C):* ISH for the expression of *Mcc*, *Lgr5* and *Pbib* (positive control) on WT and *Mcc*$^{lacZ/lacZ}$ (= *Mcc*$^{-/-}$ negative control) mouse SI sections. Endogenous *Mcc* expression is not detected on *Mcc*$^{-/-}$ tissues. *(D):* ISH for *Mcc* on transverse sections of WT and *Mcc*$^{-/-}$ SI crypts. *Mcc* expression is not detected on *Mcc*$^{-/-}$ crypts. Scale bars, 20 µm.

### Supp Figure 02

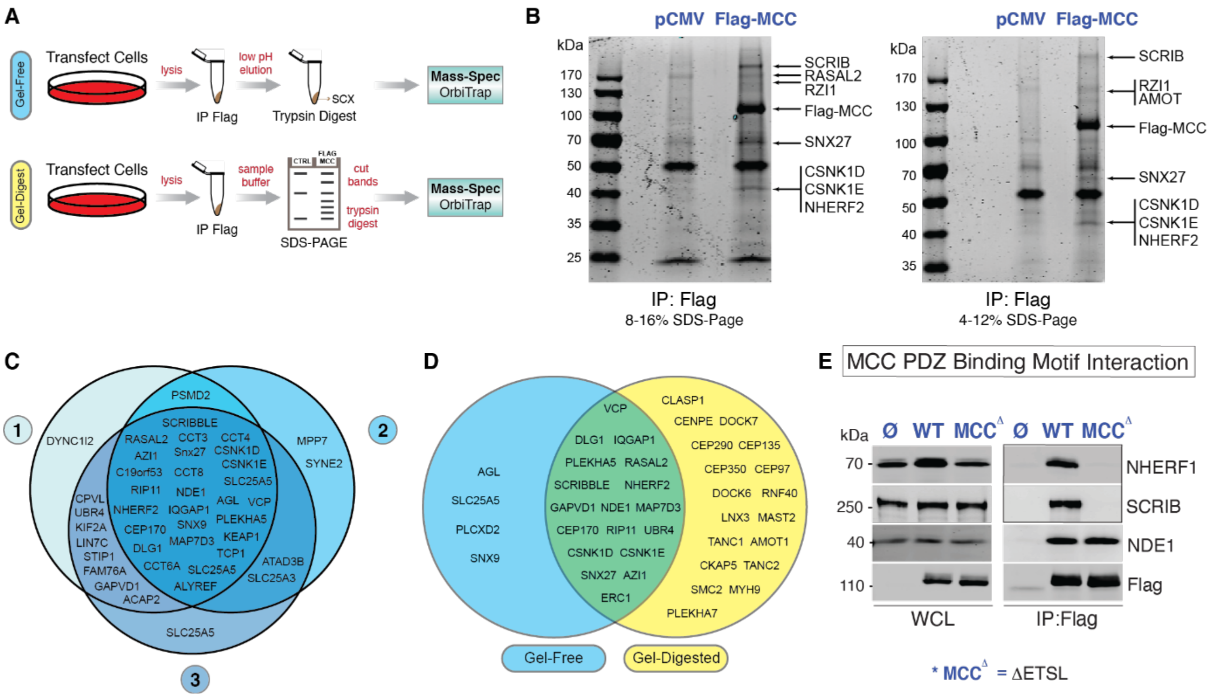


**Supplementary Figure 2. MCC interactome experiments.** *(A):* MCC interactome was captured using two approaches (Gel-free and Gel-digest). In both, N-terminal FLAG-tagged human MCC were transfected into HEK293 cells. G418-resistant cells were immunoprecipitated using anti-FLAG agarose beads. Cells stably expressing an empty vector were used as control. To capture interactions, samples were run either on an SDS-Page gel (Gel-digest approach) or eluted and separated with SCX (strong-cation exchange). SDS-Page gel samples were run against an empty vector control (*pCMV* empty) and bands present in the FLAG-MCC lane *(B)* were extracted. Samples from both approaches were subjected to mass spectrometry (OrbiTrap) for protein identification and quantification. *(C):* Venn diagram analysis of three replicates (Gel-free approach) revealed significant overlap in MCC interactors. *(D):* Venn diagram analysis of the two mass-spectrometry approaches (Gel-free and Gel-digested) show a significant number of interactions present in both samples. *(E):* Truncation of the PDZ binding motif (PBM) (-ETSL) of MCC (MCCΔ) disrupts interaction to NHERF1 and SCRIB, but not to NDE1 in HEK293 cells. WCL, whole cell lysate. N = 3. IP: Immunoprecipitation. IB: Immunoblotting.

**Supp Figure 03**

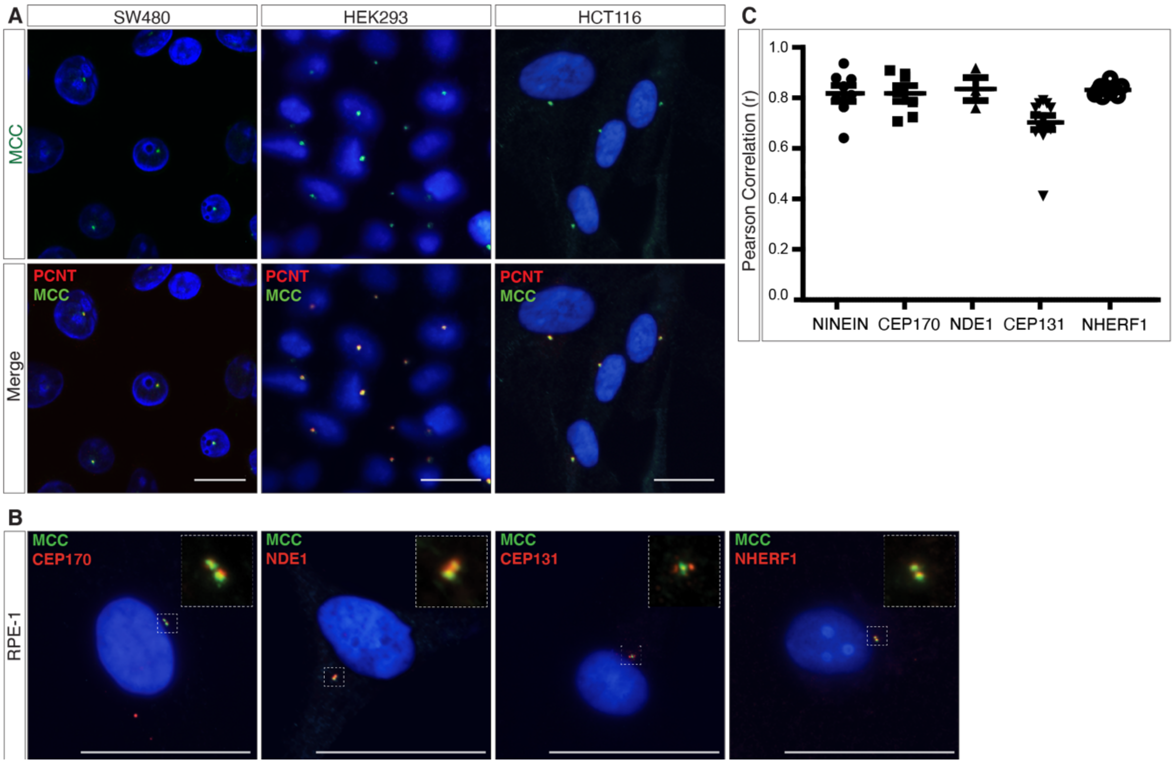


**Supplementary Figure 3. MCC colocalizes with well-characterized centrosomal proteins in multiple human cell lines.** *(A):* Immunofluorescence (IF) shows colocalization of MCC and PERICENTRIN (PCNT) at the centrosome in SW480, HEK293, and HCT116 cells. Scale bars, 50 μm. *(B):* IF in RPE-1 cells shows colocalization of MCC and CEP170, NDE1, CEP131, and NHERF1. *(C):* Pearson correlation coefficient (r) reveals strong positive linear correlation between the signal positions of MCC and several of its interacting partners at the centrosome in RPE-1 cells.

### Supp Figure 04

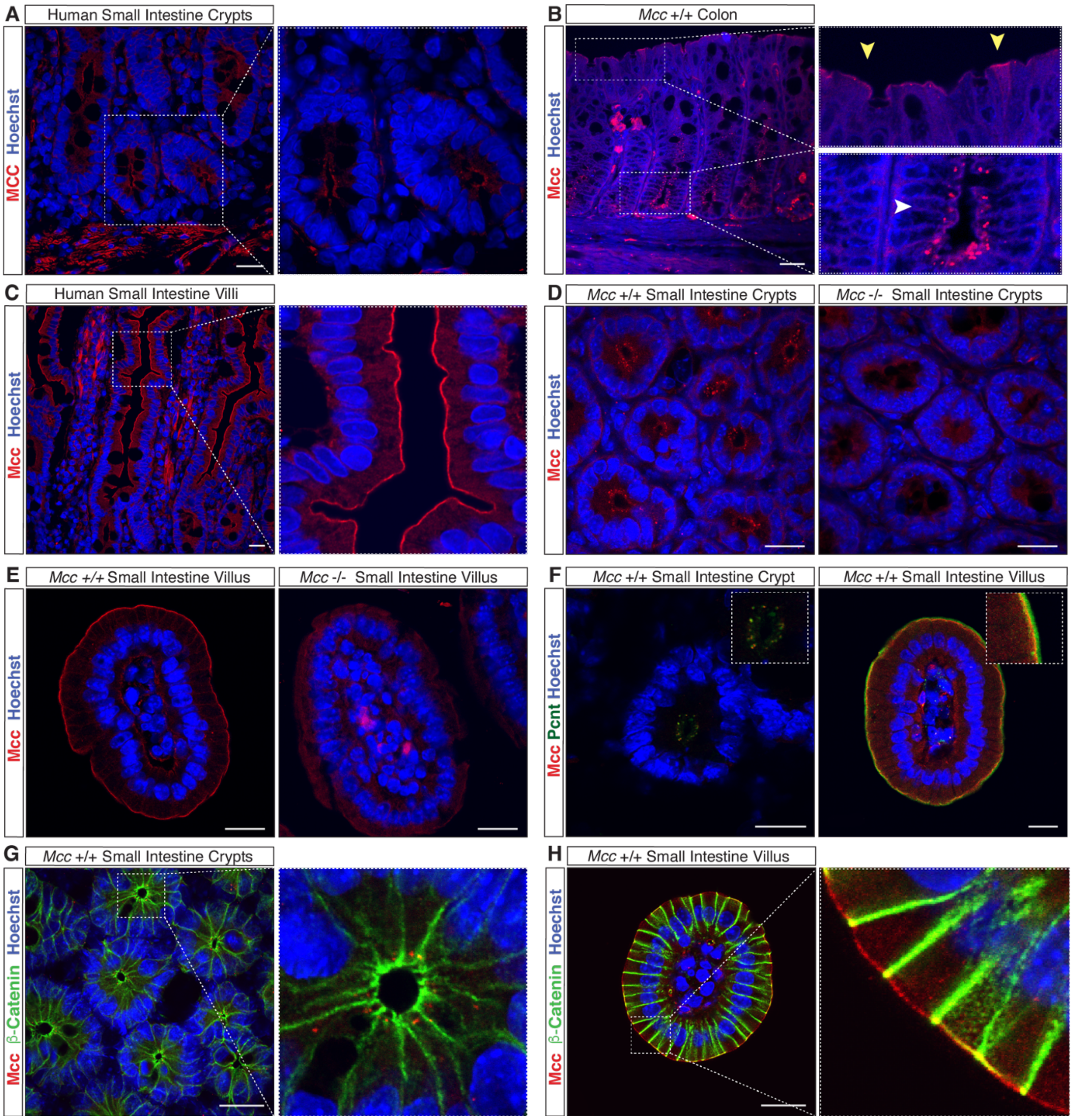


**Supplementary Figure 4. MCC antibody specificity control for immunofluorescence (IF) in the intestine.** *(A):* IF for MCC on sections of human SI shows MCC localizing at the centrosome in crypt cells. *(B):* IF for Mcc localization on WT (wild-type) mouse colonic epithelium reveals Mcc at the centrosome in crypt cells (white arrowhead) and at the apical membrane of differentiated cells (yellow arrowheads). *(C):* IF for MCC on sections of human SI shows MCC localizing at the apical membrane of differentiated cells in villi. *(D-E):* IF for Mcc localization on WT and $Mcc^{lacZ/lacZ}$ (= $Mcc^{-/-}$ negative control) transverse sections of the mouse small intestine (SI). *(D):* Mcc localizes to the centrosome (punctate staining) in crypts and *(E)* at the apical membrane of differentiated cells in the villus units. $Mcc^{-/-}$ sections show absence of Mcc signal in crypts *(D)* and villi *(E)*. *(F):* IF for Mcc and Pcnt shows colocalization at the centrosome in crypt cells and at the apical membrane of villus cells (transverse sections). *(G-H):* IF for Mcc and β-catenin on a WT transverse section of crypts *(G)* and villus *(H)* shows colocalization at apico-lateral cell junctions. Scale bars, 20 µm.
